# A weighted burden test using logistic regression for integrated analysis of sequence variants, copy number variants and polygenic risk score

**DOI:** 10.1101/306191

**Authors:** David Curtis

**Affiliations:** UCL Genetics Institute, UCL, Darwin Building, Gower Street London WC1E 6BT, UK; Centre for Psychiatry, Barts and the London School of Medicine and Dentistry, Charterhouse Square, London EC1M 6BQ, UK

**Keywords:** Burden test, association analysis, logistic regression

## Abstract

Previously described methods of analysis allow variants in a gene to be weighted more highly according to rarity and/or predicted function and then for the variant contributions to be summed into a gene-wise risk score which can be compared between cases and controls using a t test. However this does not allow incorporating covariates into the analysis. Schizophrenia is an example of an illness where there is evidence that different kinds of genetic variation can contribute to risk, including common variants contributing to a polygenic risk score (PRS), very rare copy number variants (CNVs) and sequence variants. A logistic regression approach has been implemented to compare the gene-wise risk scores between cases and controls while incorporating as covariates population principal components, the PRS and the presence of pathogenic CNVs and sequence variants. A likelihood ratio test is performed comparing the likelihoods of logistic regression models with and without this score. The method was applied to an ethnically heterogeneous exome-sequenced sample of 6000 controls and 5000 schizophrenia cases. In the raw analysis the test statistic is inflated but inclusion of principal components satisfactorily controls for this. In this dataset the inclusion of the PRS and effect from CNVs and sequence variants had only small effects. The set of genes which are FMRP targets showed some evidence for enrichment of rare, functional variants among cases (p=0.0005). This approach can be applied to any disease in which different kinds of genetic and non-genetic risk factors make contributions to risk.

## Introduction

Variants affecting risk of disease may be individually too rare to generate statistically significant results in case-control studies and so a burden test may be performed to assess whether there is an excess among cases of particular categories of variant within a gene or set of genes, the variants typically being included based on rarity and predicted function^1,2^. Rather than exclude less rare variants, one may perform a weighted analysis in which common variants are included but are accorded less weight less than rare ones^3^. We have previously developed a method which provides weights based both on rarity and predicted functional effect^4,5^. The method performs a weighted burden analysis to test whether, in a particular gene or set of genes, variants which are rarer and/or predicted to have more severe functional effects occur more commonly in cases than controls. Each variant is assigned a weight based on its rarity and predicted function and then an overall gene-wise risk score is allocated to each subject consisting simply of the sum of the weights of the variants which are found in that subject. These scores for cases and controls are then compared by carrying out a t test. The t test is rapid to compute and statistically robust and the method performed acceptably when applied to real samples. However a t test does not incorporate information from covariates and there are two reasons why this is an important limitation. The first reason is that if the samples are ancestrally heterogeneous then artefactual results can be obtained and that using ancestry principal components as covariates might mitigate this problem. The second reason is that other measurable genetic and non-genetic factors might be known to contribute to risk of disease and that incorporating these as covariates might be expected to enhance the accuracy of the analysis. Three obvious kinds of genetic risk to consider are the polygenic risk score (PRS), the presence of known pathogenic copy number variants (CNVs) and the presence of known pathogenic sequence variants. To use schizophrenia as a concrete example, there is evidence that certain CNVs greatly increase risk but these are very rare, even among cases^6^. Likewise very rare variants causing loss of function (LOF) of a small number of genes substantially increase risk^7,8^. Statistical evidence demonstrates that rare, damaging variants in additional genes also affect risk and that these genes are concentrated in particular gene sets although the individual genes contributing to this effect are yet to be identified^9,10^. Finally, cases tend to have a higher PRS, reflecting the combined effect of many common variants, widely distributed and individually having very small effects on risk^11^. When carrying out a case-control study of exome sequence data in order to detect associations with rare, damaging variants it might be reasonable to suppose that cases with pathogenic CNVs or sequence variants might be unlikely to possess additional rare risk factors. Likewise, a case with a very low PRS might be thought more likely to possess some additional risk factor than one whose PRS is very high. Thus it seems desirable to incorporate information about different kinds of risk factor jointly when possible. Accordingly software was developed that would compare the gene-wise risk scores using logistic regression analysis so that any desired covariates could be included.

## Methods

The previously described SCOREASSOC program was modified to carry out logistic regression analysis^4,5^. It accepts as input genotypes of variants within a gene for cases and controls, with each variant assigned a weight according to its annotation as obtained using VEP, PolyPhen and SIFT^12–14^. The functional weight is then multiplied by a weight for rarity, so that rarer variants are assigned higher weights. For each subject a gene-wise risk score is derived as the sum of the variant-wise weights, each multiplied by the number of alleles of the variant which the given subject possesses. If a subject is not genotyped for a variant then they are assigned the subject-wise average score for that variant. The program was modified to accept as additional input an arbitrary number of quantitative covariates for each subject, typically population principal components, PRS and an indicator variable denoting whether or not the subject possesses a known pathogenic CNV or sequence variant. The score and covariates are entered into a standard logistic regression model with case-control status as the outcome variable and after variable normalisation the likelihood of the model is maximised using the L-BFGS quasi-newton method, implemented using the *dlib* library^15^. The contribution of different variables to risk is assessed using standard likelihood ratio tests by comparing twice the difference in maximised log likelihoods between models with and without the variables of interest. This likelihood ratio statistic is then taken as a chi-squared statistic with degrees of freedom equal to the difference between models in number of variables fitted. The coefficients for each variable can be varied to maximise the likelihood or can be fixed. For example, if it is known that a particular CNV is associated with a ten-fold increase in risk then the coefficient can be set to ln(10) to reflect this, rather than fitting it from the available dataset, which may contain fewer subjects than those used to produce the original risk estimate.

Preliminary analyses indicated that simple logistic regression could produce extreme p values if a gene had only a single very rare variant found in only one or two cases. This seems to arise because subjects with unknown genotypes are then assigned the average score for this variant. This average score is very low but non-zero while subjects who are genotyped will have scores of zero. If more cases than controls have an unknown genotype then the maximisation routine would overfit the model and would assign a very high value to the score coefficient and would produce an apparently significant likelihood ratio statistic. To address this, the ridge penalty function, consisting of the sum of the squares of the regression coefficients, was subtracted from the log likelihood. This satisfactorily prevented the artefactual extreme p values without preventing the ability to fit the model to produce the expected large coefficients for covariates such as principal components and the PRS.

The program outputs the coefficients for the fitted models along with their estimated standard errors and the results of the likelihood ratio test. When association with the gene-wise risk score alone is tested, i.e. when the two models differ only in whether or not the score is included, then the statistical significance is summarised as a signed log p value (SLP) which is the log base 10 of the p value given a positive sign if the score tends to be higher in cases and negative if it tends to be lower. For other analyses the minus log base 10 of the p value (MLP) is output. The support program for SCOREASSOC, called GENEVARASSOC, was also modified to facilitate incorporating the covariates and specifying the desired analyses. Both are implemented in C++ and can be downloaded along with documentation from the site listed below.

### Example application

The approach was applied to whole exome sequence data from the Swedish schizophrenia study, consisting of 4968 cases and 6245 controls^9^. The sequence data was downloaded as a VCF file from dbGAP (https://www.ncbi.nlm.nih.gov/gap). One aspect of special interest about this dataset is that although it was recruited in Sweden some subjects have a substantial Finnish component to their ancestry and that this applies more to cases than controls. It was analysed previously using SCOREASSOC to carry out a weighted burden test but to do this it was first necessary to remove the subjects with Finnish ancestry because otherwise some genes produced false positive results^10^. To obtain the gene-wise risk scores the same methods were used as for this previous analysis. Variants were excluded if they did not have a PASS in the Variant Call Format (VCF) information field and individual genotype calls were excluded if they had a quality score less than 30. Sites were also excluded if there were more than 10% of genotypes missing or of low quality in either cases or controls or if the heterozygote count was smaller than both homozygote counts in both cohorts. Each variant was annotated using VEP, PolyPhen and SIFT^12–14^. GENEVARASSOC was used to generate the input files for SCOREASSOC and the default weights provided with the software were used, for example consisting of 5 for a synonymous variant and 20 for a stop gained variant, except that 10 was added to the weight if the PolyPhen annotation was possibly or probably damaging and also if the SIFT annotation was deleterious. The full set of weights used is shown in Table 1. SCOREASSOC also weights rare variants more highly than common ones but because it is well-established that no common variants have a large effect on the risk of schizophrenia we excluded variants with MAF>0.01 in the cases and in the controls, so in practice weighting by rarity had negligible effect.

**Table 1.** The table shows the weight which was allocated to each type of variant according to its annotation by VEP^12^ 10 was added to this weight if the variant was annotated by Polyphen as possibly or probably damaging and 10 was added if SIFT annotated it as deleterious^13,14^.

| VEP annotation | Weight |
| --- | --- |
| intergenic_variant | 1 |
| feature_truncation | 3 |
| regulatory_region_variant | 3 |
| feature_elongation | 3 |
| regulatory_region_amplification | 3 |
| regulatory_region_ablation | 3 |
| TF_binding_site_variant | 3 |
| TFBS_amplification | 3 |
| TFBS_ablation | 3 |
| downstream_gene_variant | 3 |
| upstream_gene_variant | 3 |
| non_coding_transcript_variant | 3 |
| NMD_transcript_variant | 3 |
| intron_variant | 3 |
| non_coding_transcript_exon_variant | 3 |
| 3_prime_UTR_variant | 5 |
| 5_prime_UTR_variant | 5 |
| mature_miRNA_variant | 5 |
| coding_sequence_variant | 5 |
| synonymous_variant | 5 |
| stop_retained_variant | 5 |
| incomplete_terminal_codon_variant | 5 |
| splice_region_variant | 5 |
| protein_altering_variant | 10 |
| missense_variant | 10 |
| inframe_deletion | 15 |
| inframe_insertion | 15 |
| transcript_amplification | 15 |
| start_lost | 15 |
| stop_lost | 15 |
| frameshift_variant | 20 |
| stop_gained | 20 |
| splice_donor_variant | 20 |
| splice_acceptor_variant | 20 |
| transcript_ablation | 20 |

To obtain population principal components, the genotypes were thinned to include only variants present on the Illumina Infinium OmniExpress-24 v1.2 BeadChip (http://emea.support.illumina.com/downloads/infinium-omniexpress-24-v1-2-product-files.html) and then version 1.09beta of *plink* (https://www.cog-genomics.org/plink2) was run with the options --*pca header tabs --make-rel*^16–18^. In order to obtain a PRS for schizophrenia, the file called *scz2.prs.txt.gz,* containing ORs and p values for 102,636 SNPs, was downloaded from the Psychiatric Genetics Consortium (PGC) website (*www.med.unc.edu/pgc/results-and-downloads*). This training set was produced as part of the previously reported PGC2 schizophrenia GWAS^n^. SNPs with p value < 0.05 were selected and their log(OR) summed over sample genotypes using the *--score* function of *plink 1.09beta* in order to produce a PRS for each subject^16–18^.

Attempts were made to use allele depth information in the VCF file to call all the CNVs with odds ratio (OR) reported to be greater than 9 as listed in Table 1 of a recent study of over 40,000 subjects^6^. A deletion was called if there was a relatively low read depth over the region (compared to the expected depth across subjects for each variant and across variants for each subject) and if there were very few heterozygote calls. A duplication was called if there was a relatively high number of reads and if the heterozygote calls tended to occur with allele ratios of 2:1 or 1:2 instead of 1:1. To accomplish this, a two-stage process was used, consisting of an automated short-listing of subjects followed by visual inspection of detailed results for these short-listed subjects. To carry out the short-listing process, for each variant a likelihood for the observed allele depths was calculated according to the copy number being 0 (deletion), 1 or 2 (duplication) and from this a log likelihood ratio statistic (LRS) favouring either deletion or duplication was derived. Then for each subject a t test was used to compare the LRS of variants within the CNV region to those outside it and another t test was used to compare the overall depth of variants inside and outside the CNV region. The short-listed subjects consisted of the 30 subjects having test results most strongly suggesting a deletion or duplication, along with 30 subjects having the lowest or highest average difference in depths, producing a possible maximum of 120 subjects. For these subjects, graphs were produced showing the LRS and moving average LRS along with the allele ratios and moving average depths. CNV calls were made blind to phenotype based on visual inspection of these graphs and a judgement as to whether the overall pattern seemed consistent with the presence of a deletion or duplication. It can be difficult to detect CNVs from exome-sequence VCF files because some regions will not be covered at all and because depth information is only provided for positions where a variant allele is observed. There was insufficient information to call the CNV at 9:831690-959090 and no subjects were called with a CNV at 3:197230000-198840000, although the latter is very rare and it may be that it was not present. The number of calls that were made in cases and controls is shown in Table 2. Although there is a marked excess of CNV calls among cases at 16:29560000-30110000 and 22:17400000-19750000, this is not the case for the other locations and it assumed that many of these calls may be erroneous. The intent of the current study is simply to demonstrate the feasibility of the analytic approach, so errors in the CNV calls are not regarded as especially problematic. However because of the unreliability of the calls it was decided not impose a fixed effect for the CNVs but to fit the effect size as observed in this dataset. For brevity, these CNVs having large effects of risk, along with LOF sequence variants having large effects on risk, will be referred to as pathogenic.

**Table 2.** Counts of CNV calls in cases and controls for those CNVs with OR>9 in Table 1 from a recent study of CNVs in schizophrenia^6^. Attempts were made to call the CNVs from the VCF file, which only contains depth information about locations within exons where there is allelic variation. There was insufficient information to call the CNV at 9:831690-959090 and it is expected that some of the other calls are erroneous.

| Position | Pathogenic CNV | Controls, N=6245 | Cases, N=4968 |
| --- | --- | --- | --- |
| 2:50000992-51113178 | del | 6 | 8 |
| 3:197230000-198840000 | del | 0 | 0 |
| 7:72380000-73780000 | dup | 0 | 3 |
| 9:831690-959090 | del/dup | - | - |
| 8:100094670-100958984 | del | 9 | 9 |
| 15:28920000-30270000 | del | 21 | 18 |
| 16:28730000-28960000 | del | 1 | 4 |
| 16:29560000-30110000 | dup | 2 | 14 |
| 22:17400000-19750000 | del | 0 | 9 |
| Any pathogenic CNV |  | 38 | 65 |

Based on previous findings, it was decide to regard LOF sequence variants in SETD1A, RBM12 or NRXN1 as pathogenic^6–8^,^19^,^20^. No subjects had a LOF variant in NRXN1 or RBM12. However two cases had LOF variants in SETD1A, with one having a stop gained variant and the other a splice acceptor variant.

Likelihood ratio tests were carried out to assess the association of the principal components with caseness and then, using the principal components as covariates, to assess the association of the PRS and of the pathogenic CNVs and sequence variants.

For each gene, the following analyses were carried out to test the association of the gene-wise risk score with caseness:

A t test comparing scores in cases and controls.

Logistic regression analysis of the scores with no covariates.

Analysis of the scores including principal components as covariates.

Analysis of the scores including principal components and PRS as covariates.

Analysis of the scores including principal components and pathogenic variants as covariates.

Analysis of the scores including principal components, PRS and pathogenic variants as covariates.

In order to test whether particular sets of genes demonstrated association with caseness, the gene-wise scores for all the genes in a set were summed to produce a set-wise risk score using the PATHWAYASSOC program^5^. Then the same analyses as listed above were carried out using the set-wise risk scores. Two lists of gene sets were used. The first list consisted of the 31 gene sets which had been previously tested for an enrichment for damaging or disrupting ultrarare variants (dURVs) in cases in this dataset^9^. The second list consisted of the 1454 “all GO gene sets, gene symbols” pathways downloaded from the Molecular Signatures Database at http://www.broadinstitute.org/gsea/msigdb/collections.jsp (Subramanian, Tamayo et al. 2005).

Results were managed and graphs produced using R^21^.

## Results

Carrying out the logistic regression analyses is notably slower than performing a t test and to carry out all the analyses takes a few minutes for each gene, so that analyses of all 22021 genes took a couple of days to complete on a computer cluster.

The likelihood ratio test comparing models with or without the first 20 population principal components showed that ancestry was strongly associated with caseness (X^2^=374, 20 df, MLP>35). Using these principal components as covariates, both the PRS (X^2^=156, 1 df, MLP=35) and the presence of a pathogenic CNV or sequence variant (X^2^=39.6, 8df, MLP=5.4), were also associated with caseness.

Figure 1 shows the QQ plots obtained for the different gene-wise analyses. The simple t test comparing gene-wise risk scores has a clear excess of positive SLPs above the chance expectation. The most extreme of these is for COMT, SLP=7.4. As discussed previously, this gene-wise result is largely driven by SNP rs6267 which is heterozygous in 51/6242 controls and 94/4962 cases (OR=2.3, p=8*10^-7^) and this reflects the fact that variant is much commoner in Finns than in non-Finnish Europeans^10^. One gene, CDCA8, has an extremely negative result and this reflects the fact that more controls than cases have a number of very rare variants with high functional weights. The results for the logistic regression analysis of the gene-wise risk scores alone, shown in Figure 1b, are very similar to those obtained from the t test. However when the population principal components are included, as shown in Figure 1c, the results conform almost exactly to what would be expected by chance. The SLP for CDCA8 remains somewhat low at −5.6 but this does not exceed the threshold for significance using a Bonferroni correction for the number of genes tested. Including the PRS, CNVs and SETD1A variants as covariates, as shown in Figures 1d, 1e and 1f, does not have a large impact on the overall distribution of the SLPs.

**Figure 1.**
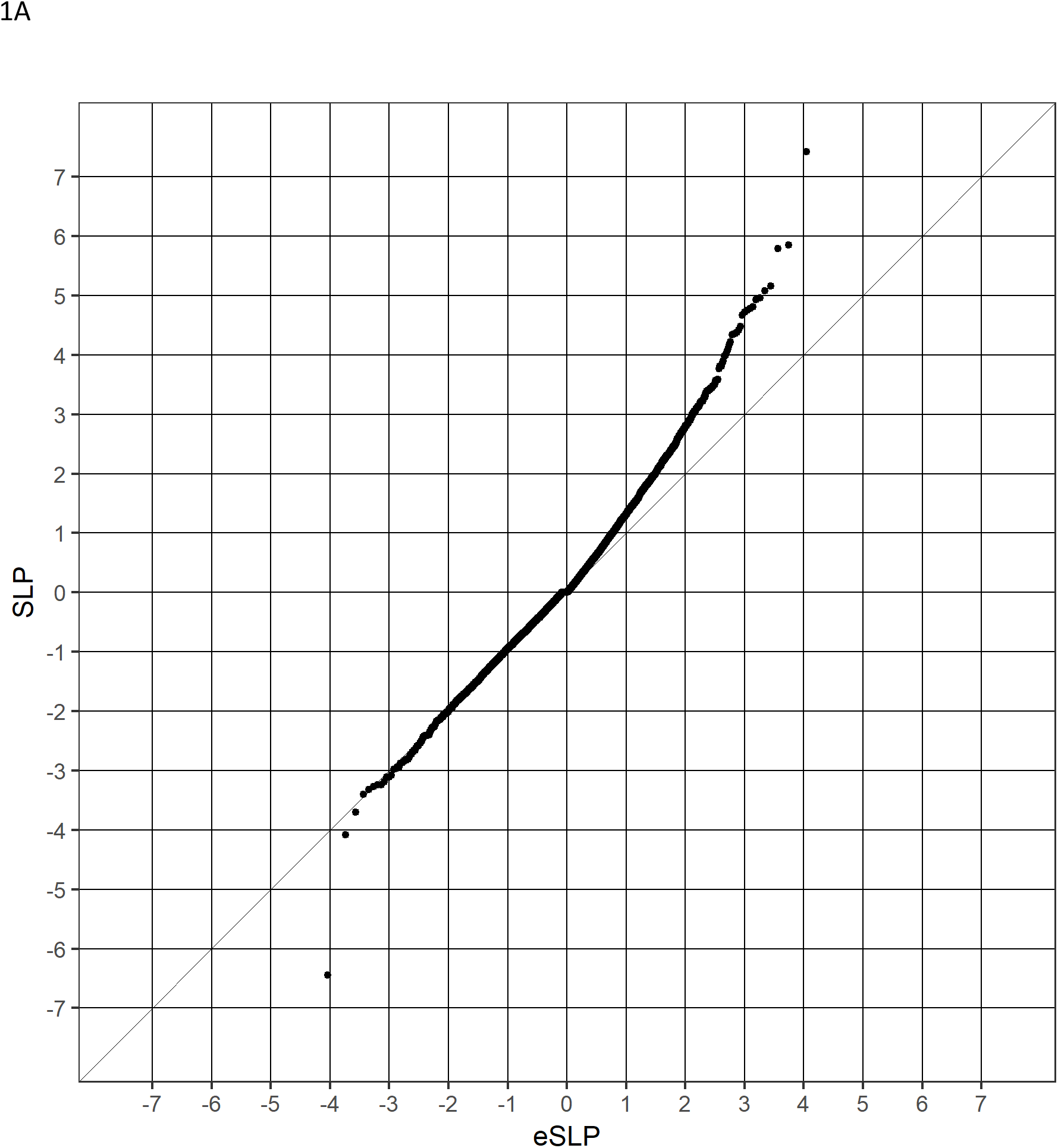
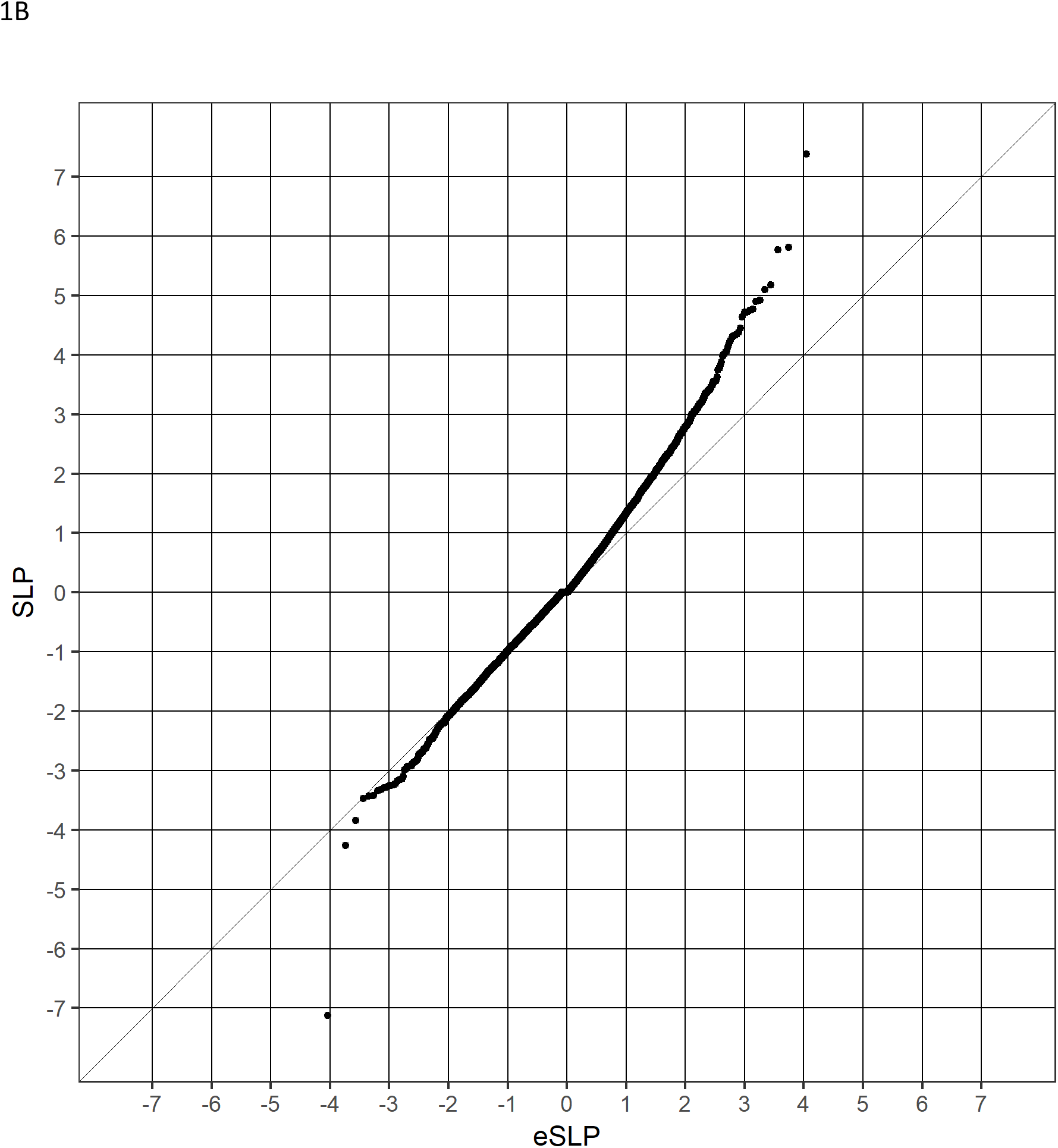
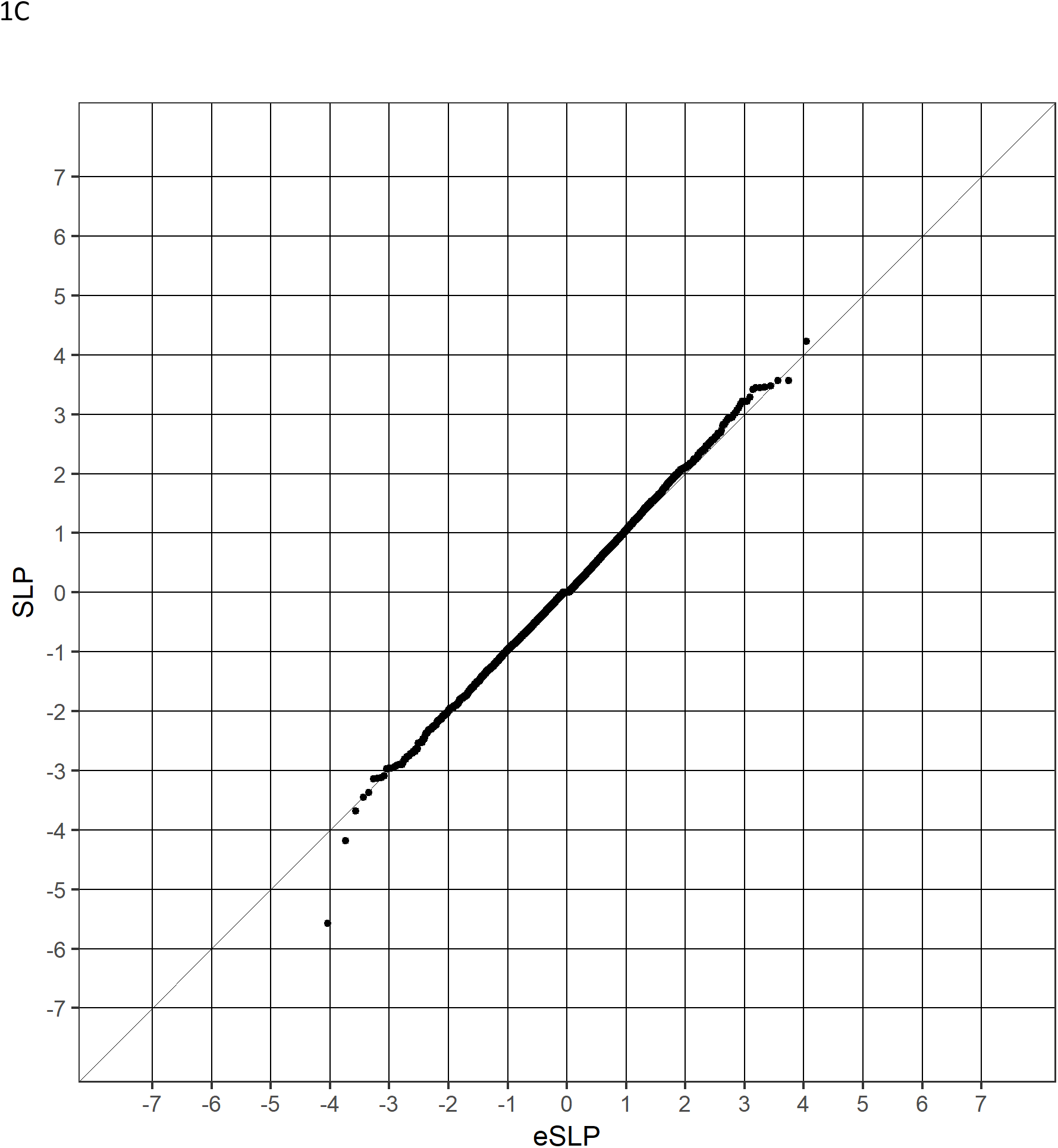
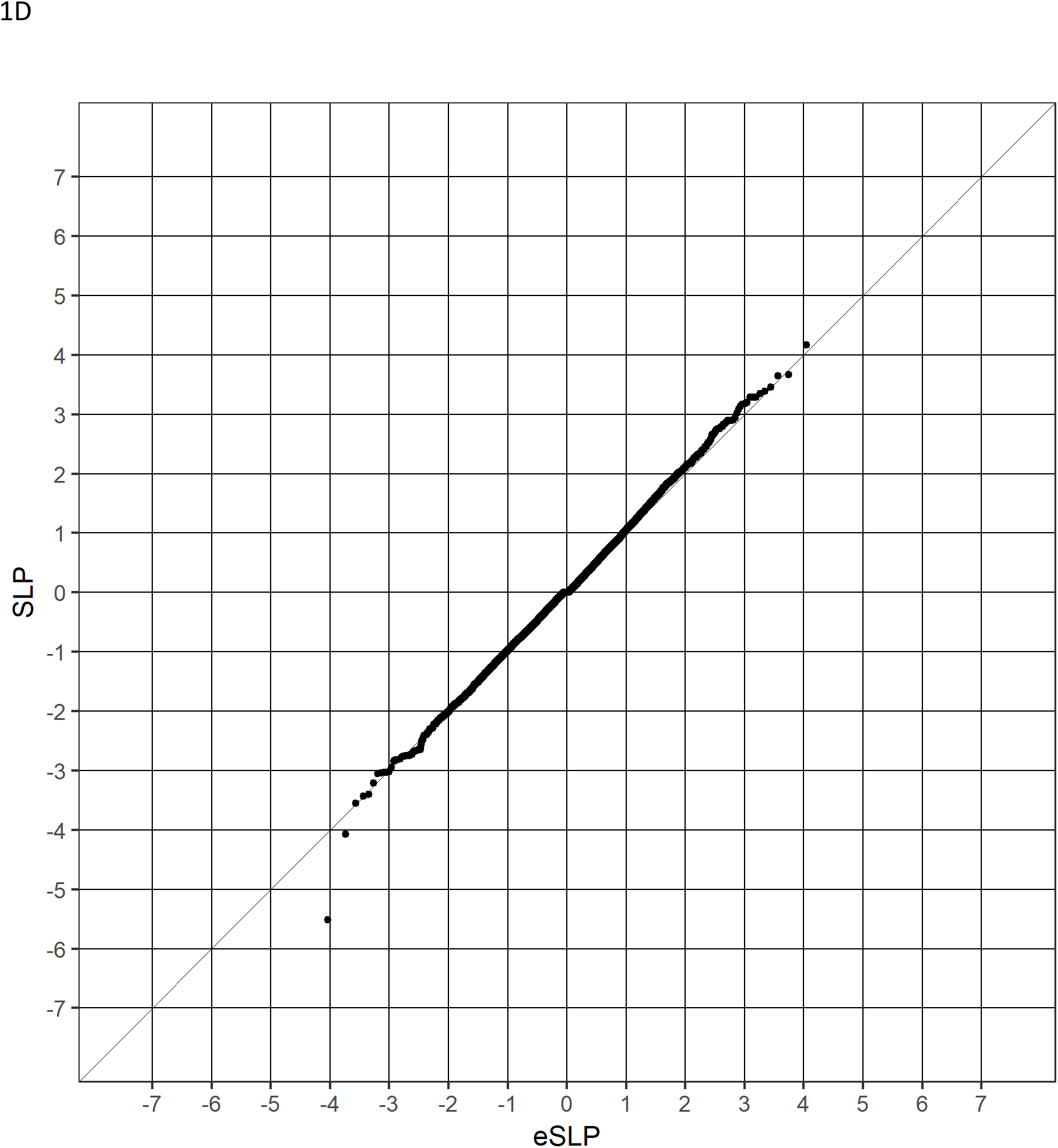
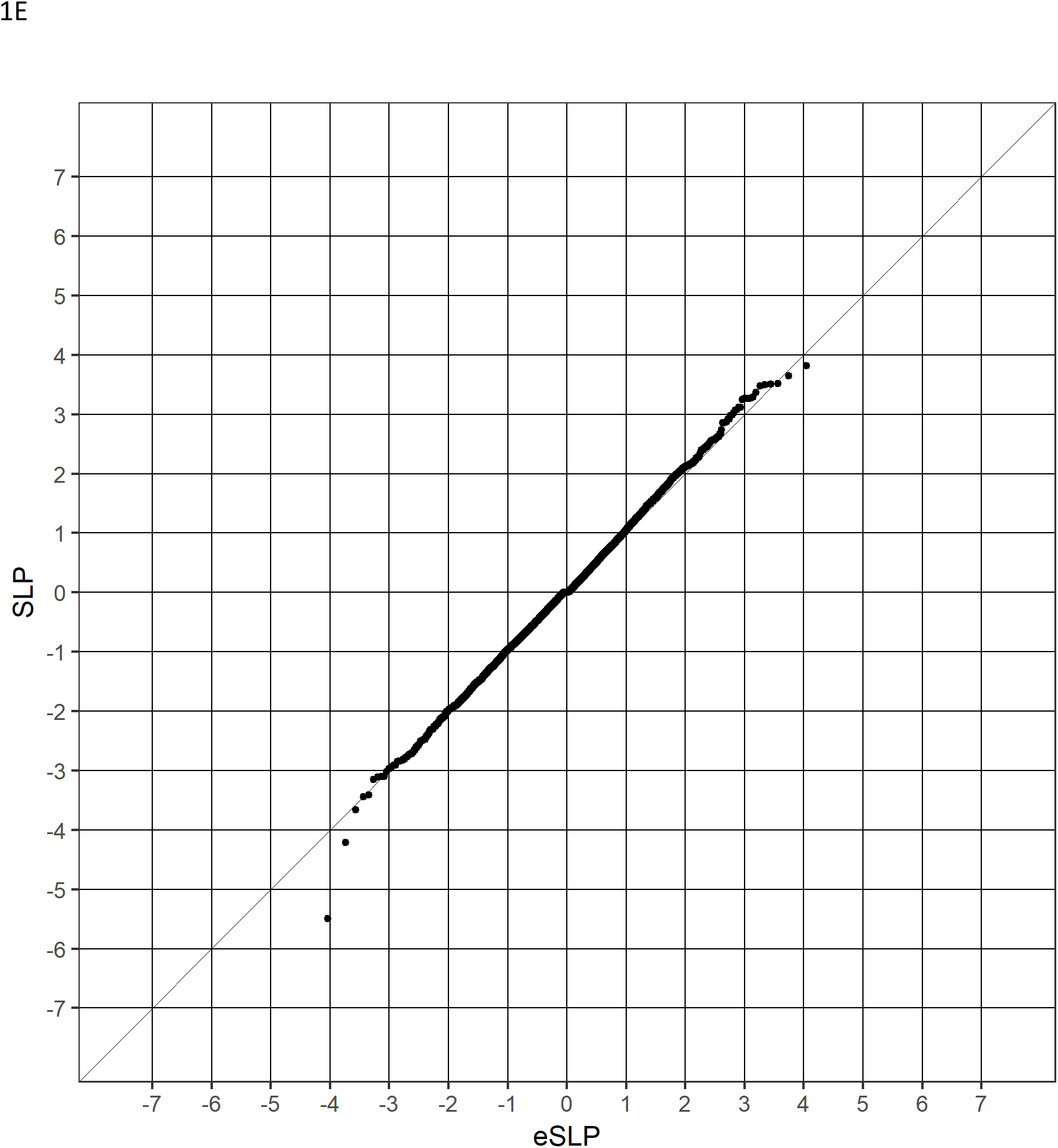
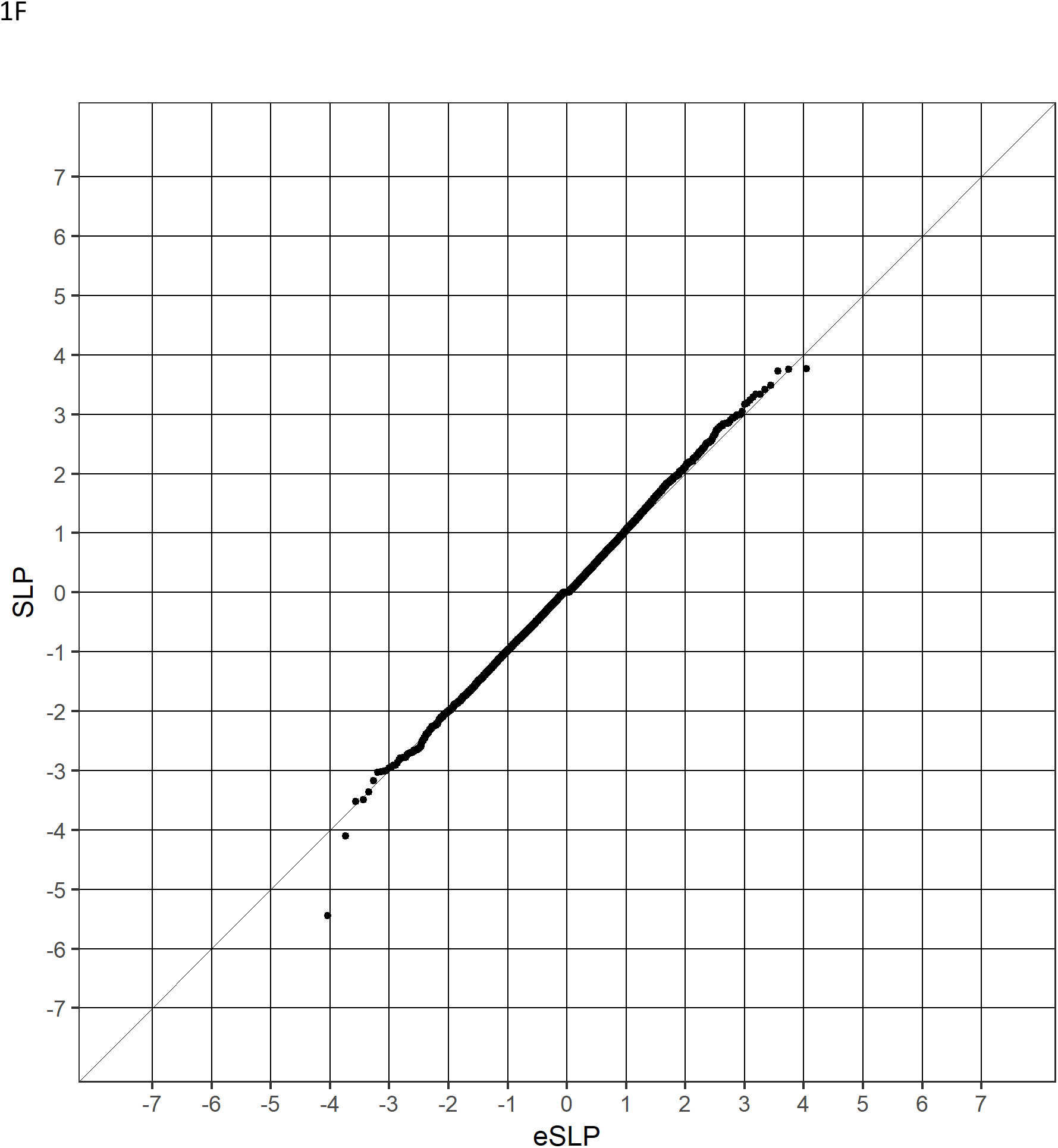
QQ plots of SLPs obtained for 22021 genes using different methods of analysis. 1A Results from t tests comparing gene-wise risk scores between cases and controls. Results for Figures 1B to 1F use logistic regression analysis of gene-wise risk scores with caseness as outcome. Analyses include the following covariates. 1B No covariates. 1C 20 population principal components. 1D 20 population principal components and PRS. 1E 20 population principal components and pathogenic CNV or sequence variants. 1E 20 population principal components, PRS and pathogenic CNV or sequence variants.

Examining the SLPs for individual genes showed that, as can be seen from the QQ plots, including the population principal components as covariates could have a large impact. The largest effect was seen with COMT, where the SLP was reduced by 4.7, and there were an additional 6 genes for which the absolute value of the SLP reduced by more than 3 and in all there were 59 genes where the magnitude of the SLP reduced by 2 or more. For all but 2 of these the SLP was positive. There were other genes where the magnitude of the SLP was increased, though to a lesser extent. For only one gene, CRISP3, did this change exceed 2 and there were 24 others for which the change was 1 or greater. All these genes had a negative SLP. Thus, when including the principal components had a large effect, the effect was generally to make positive SLPs smaller and negative SLPs larger. Although there were large effects for some genes, overall the effects were fairly small and evenly balanced, so that across all genes the mean SLP was reduced by 0.088 and the mean of the absolute value of the SLP was reduced by 0.059.

Including the PRS as a covariate had only small effects on the results. The largest change in SLP was only 0.6. Considering those genes which had the largest change in the absolute value of the SLP, there was a fairly equal distribution between those with positive and negative SLPs. The average effects across all genes were almost perfectly balanced, with the mean SLP changing by only −0.00078 and the mean absolute value of the SLP changing by 0.00016.

Including the pathogenic variants as covariates also had only small effects. For two genes the negative SLPs decreased by 0.95 and 0.87 but in no other gene did the SLP change by more than 0.51. In spite of the small magnitude of the changes, there was a very striking imbalance in the way they were distributed. Out of the 200 genes with the largest increase in the absolute value of the SLP 197 had a negative SLP while out of the 200 genes with largest decrease in the absolute value of the SLP 199 had a positive SLP. Thus, in the analyses for which including the pathogenic variants had the largest effect, doing so tended to reduce positive SLPs and increase negative SLPs. This imbalance was only notable for the analyses with the largest effects and overall the average effect was small so that including the pathogenic CNVs changed the mean SLP by 0.00018 and the mean absolute value of the SLP by 0.0013.

As in previous studies of this dataset, none of the individual gene-wise tests is statistically significant at the genome-wide level. Table 3 shows genes with SLP of 3 or higher from the analysis including all covariates.

**Table 3.** Table showing gene-wise risk score SLPs for genes achieving SLP greater or equal to 3 in the analysis including all covariates. (PCs = population principal components, PRS = schizophrenia polygenic risk score, CNV + LOF = pathogenic copy number variant or loss of function variant.)

| Gene symbol | Gene name | Analysis method |  |  |  |  |  |
| --- | --- | --- | --- | --- | --- | --- | --- |
|  |  | t test | Logistic regression using listed covariates |  |  |  |  |
|  |  |  | None | PCs | PCs,<br>PRS | PCs,<br>CNV +<br>LOF | PCs,<br>PRS,<br>CNV +<br>LOF |
| LY9 | T-lymphocyte surface<br>antigen Ly-9 | 4.7 | 4.6 | 3.4 | 3.7 | 3.5 | 3.8 |
| SNORA62 | Small Nucleolar RNA,<br>H/ACA Box 62 | 3.4 | 4.2 | 4.2 | 4.2 | 3.8 | 3.8 |
| ADAMTSL1 | ADAMTS-like protein<br>1 | 4.4 | 4.4 | 3.6 | 3.7 | 3.7 | 3.7 |
| RNU5D-1 | RNA, U5D Small<br>Nuclear 1 | 4.4 | 4.4 | 3.5 | 3.5 | 3.5 | 3.5 |
| PITPNA | Phosphatidylinositol<br>transfer protein alpha<br>isoform | 3.0 | 3.1 | 3.5 | 3.4 | 3.5 | 3.4 |
| LOC100133091 | Uncharacterized<br>LOC100133091 | 3.2 | 3.3 | 3.2 | 3.3 | 3.3 | 3.3 |
| POMZP3 | POM121 and ZP3<br>fusion protein | 3.2 | 3.3 | 3.2 | 3.3 | 3.3 | 3.3 |
| LOC101927640 | Uncharacterized<br>LOC101927640 | 2.5 | 2.5 | 3.3 | 3.3 | 3.3 | 3.3 |
| ALG6 | Dolichyl<br>pyrophosphate<br>Man9GlcNAc2 alpha-1,3-glucosyltransferase | 3.4 | 3.6 | 3.5 | 3.2 | 3.5 | 3.2 |
| KLHL11 | Kelch-like protein 11 | 4.4 | 4.3 | 3.1 | 3.2 | 3.1 | 3.2 |
| PLIN3 | Perilipin-3 | 4.8 | 4.7 | 3.5 | 3.4 | 3.3 | 3.2 |
| HNRNPA1L2 | Heterogeneous<br>nuclear<br>ribonucleoprotein A1-like 2 | 5.1 | 5.1 | 3.2 | 3.1 | 3.1 | 3.1 |
| OR9Q2 | Olfactory receptor<br>9Q2 | 2.9 | 2.9 | 2.9 | 2.9 | 3.0 | 3.0 |

The results of the gene set analyses applied to the sets used in the previous analyses of this dataset are shown in Table 4. It can be seen that many sets have a high SLP before including the PCs as covariates, demonstrating that one may be at risk of obtaining false positive results if this is not done.. When all the covariates are included the highest SLP achieved is for the set consisting of FMRP targets, with SLP=3.3, equivalent to p=0.0005.

**Table 4.**
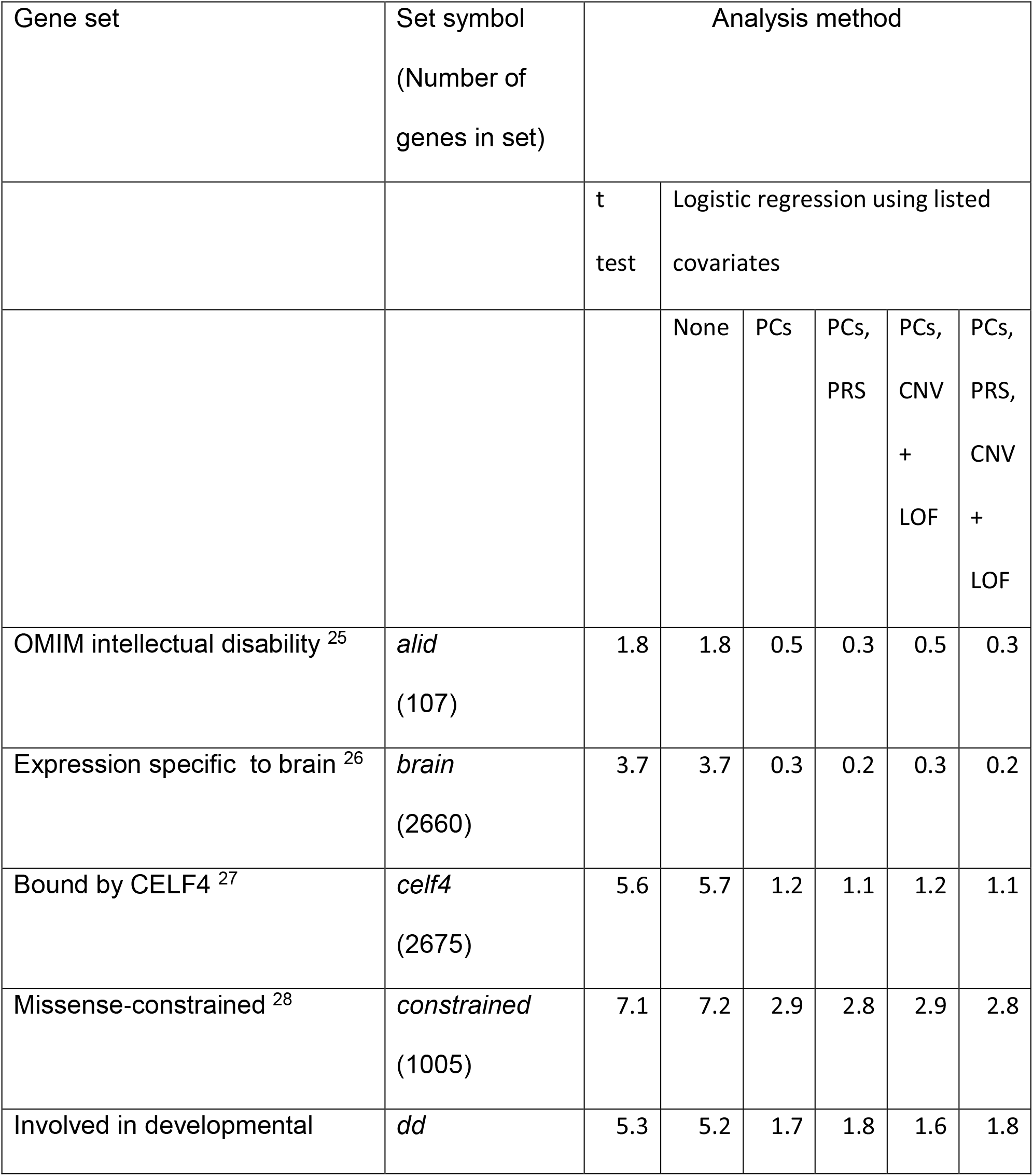
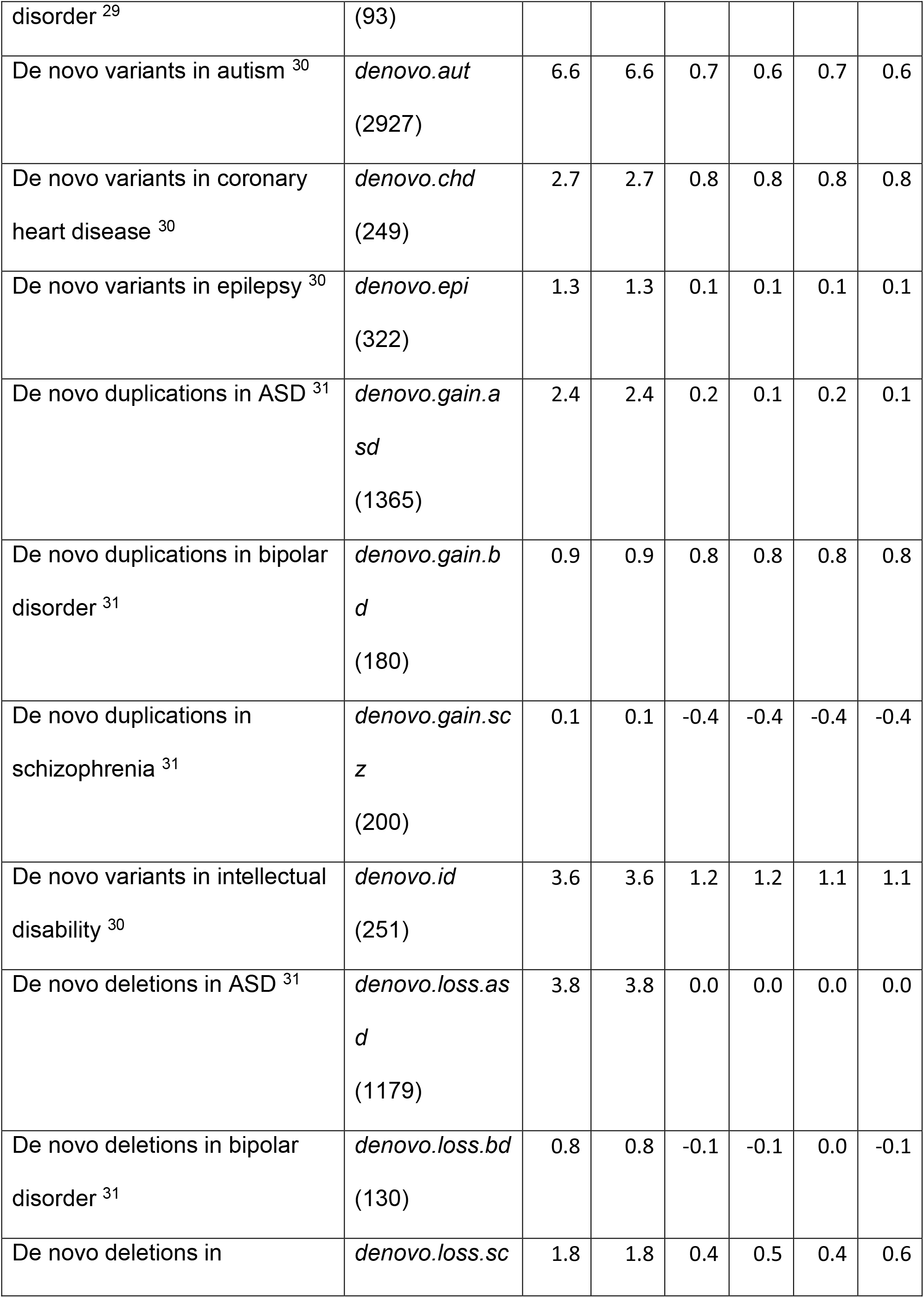
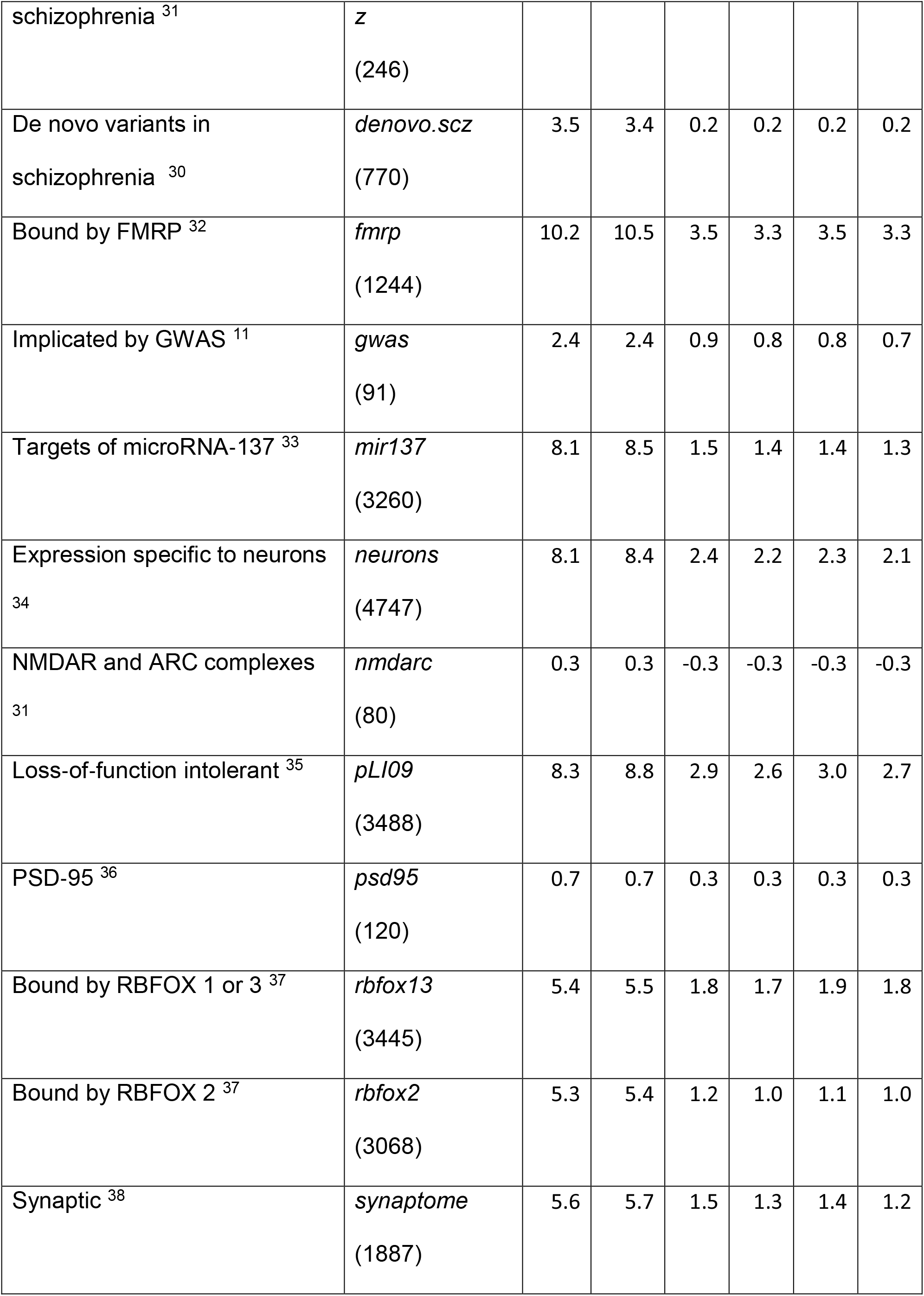
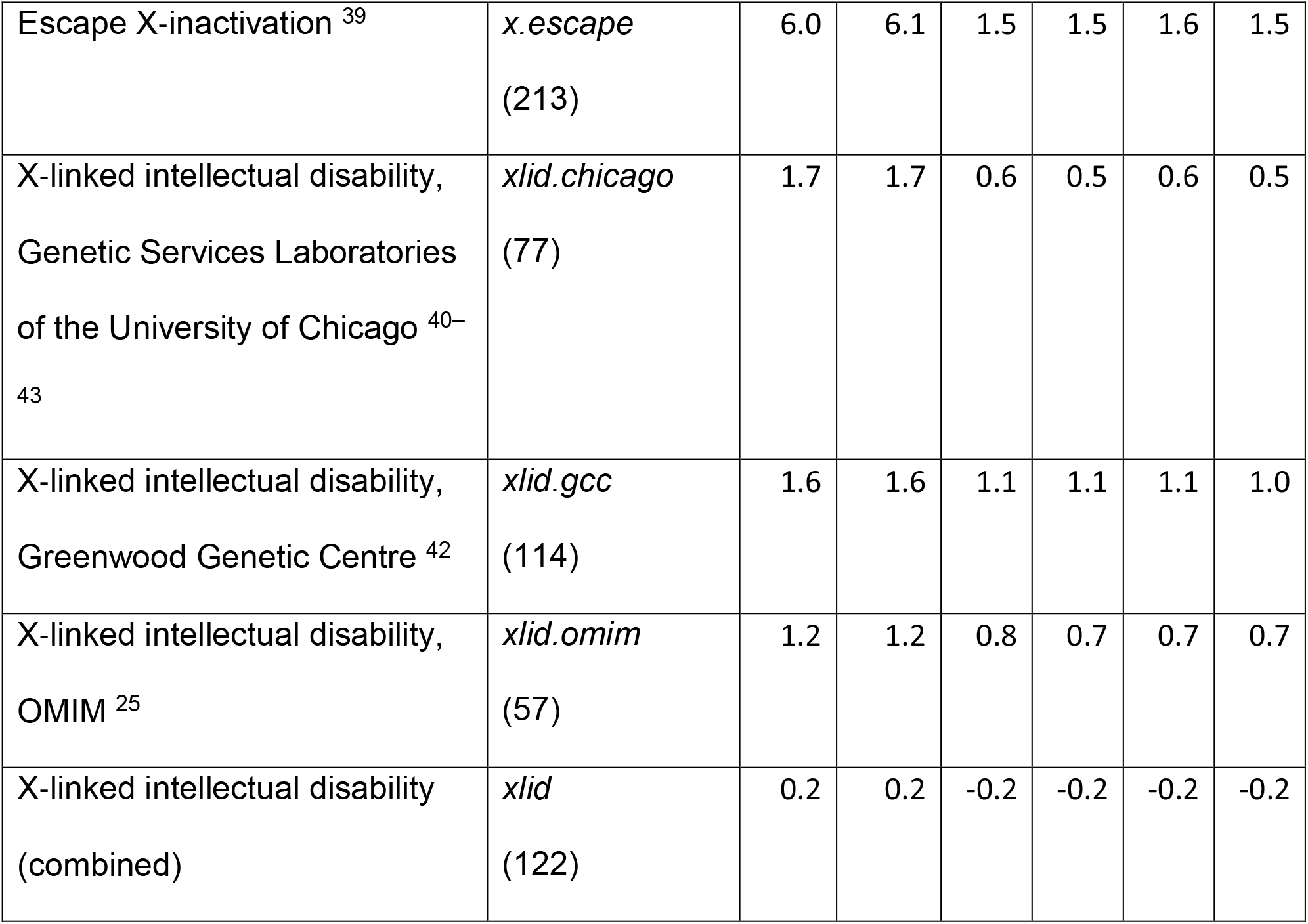
Table showing results for the gene sets used in previous analyses^9,10^.

**Table 5.**
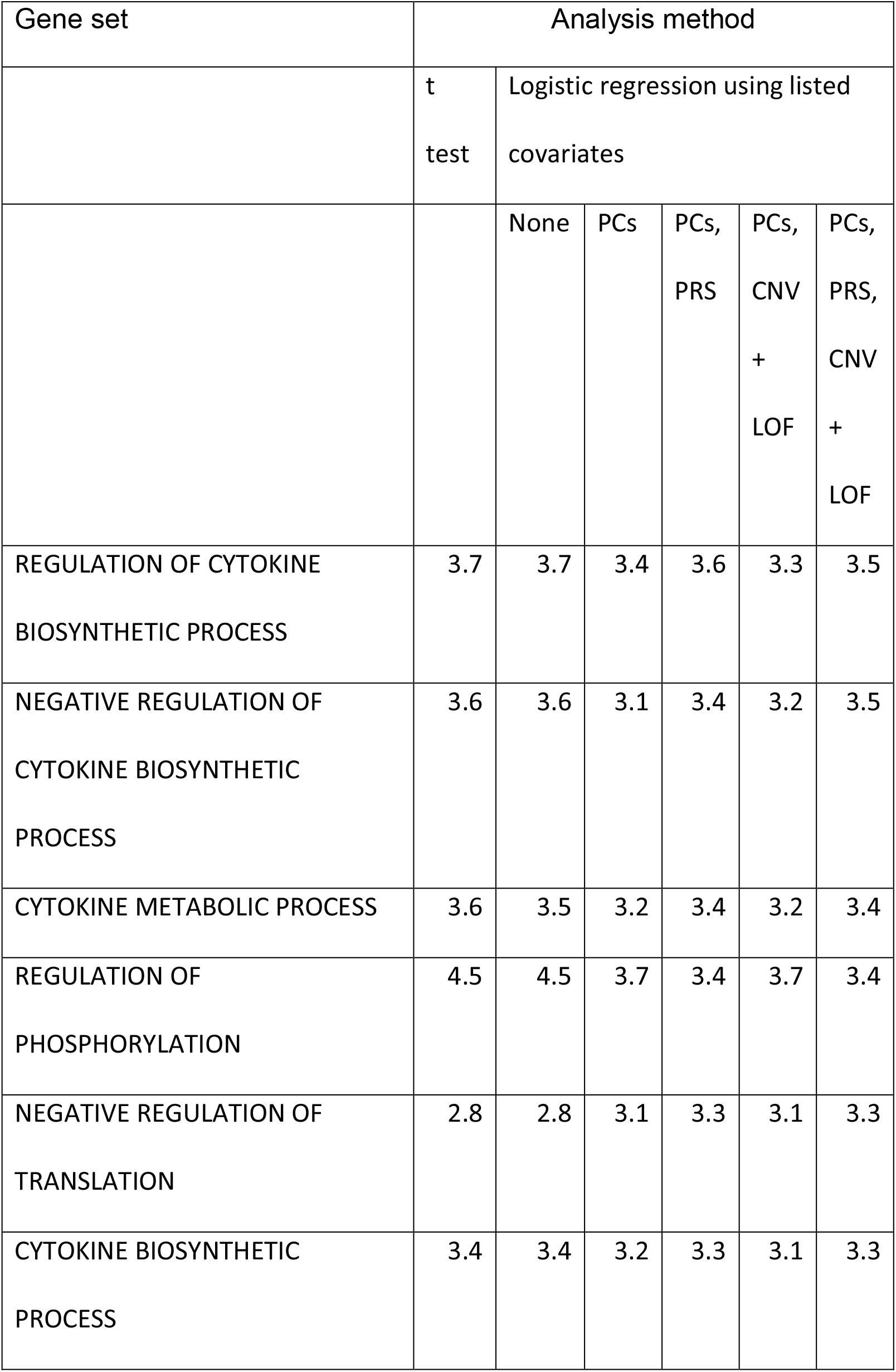
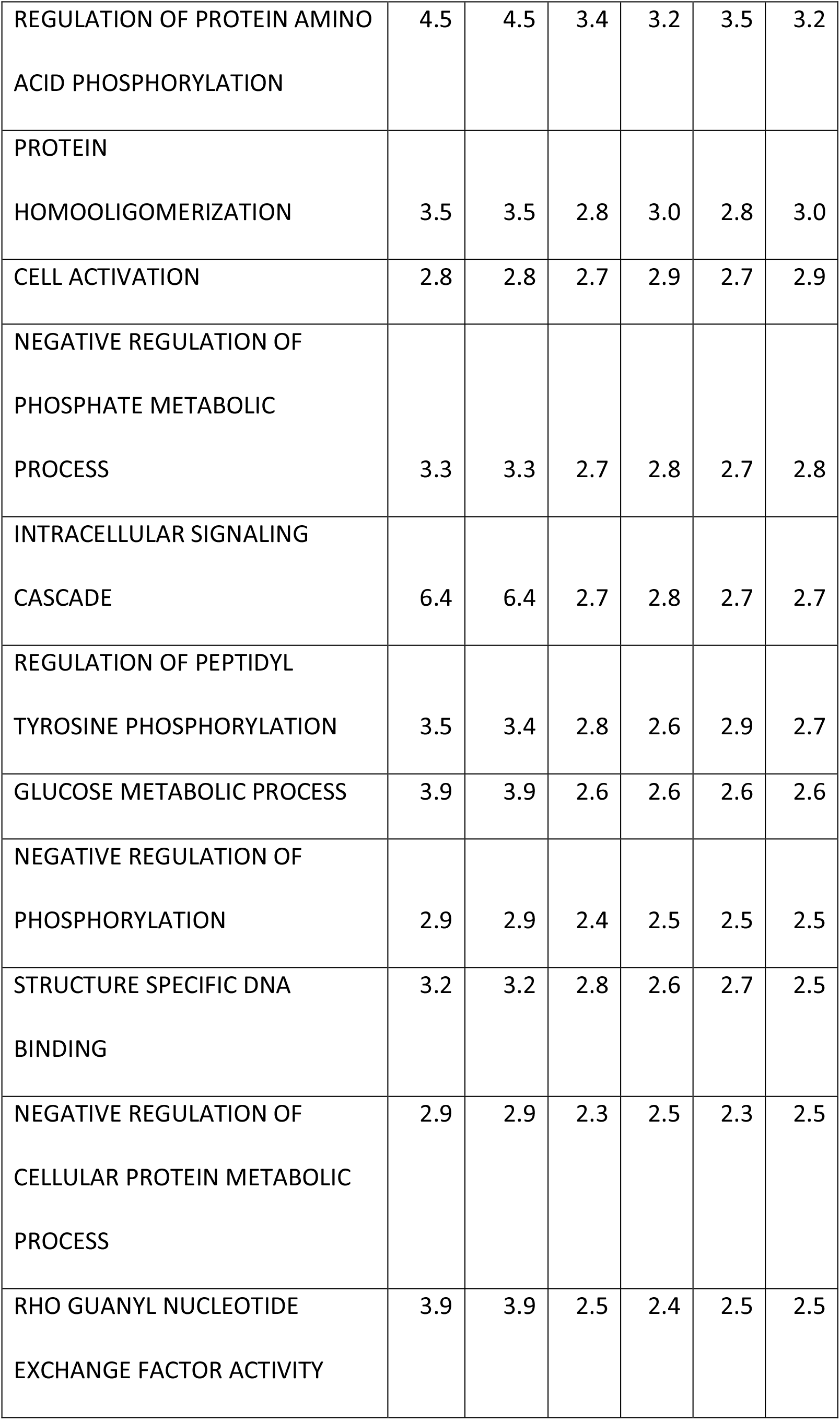
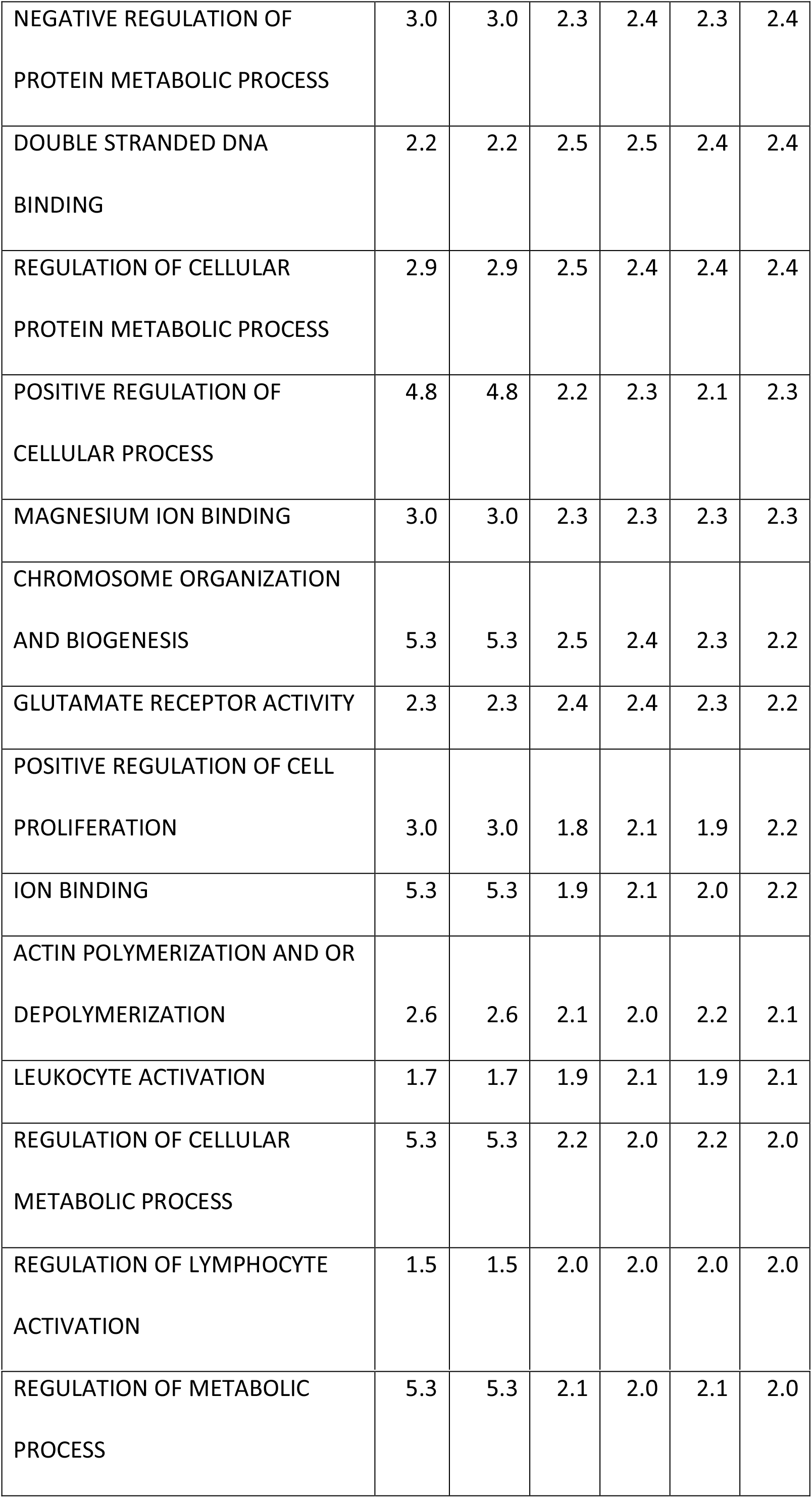
Table showing GO gene sets achieving SLP greater or equal to 2.

Figure 2 shows a QQ plot of the SLPs obtained for the GO gene sets against the expectation if the sets were independent. and it can be seen that they conform fairly closely to this distribution although with some excess of positive SLPs. In fact, the sets are not independent and genes overlap between sets so one could argue that the SLPs are somewhat higher than would be expected by chance but there is certainly no strong evidence to implicate specific sets. Table 4 shows those sets achieving an SLP of 2 or more in the full analysis including all covariates.

**Figure 2.**
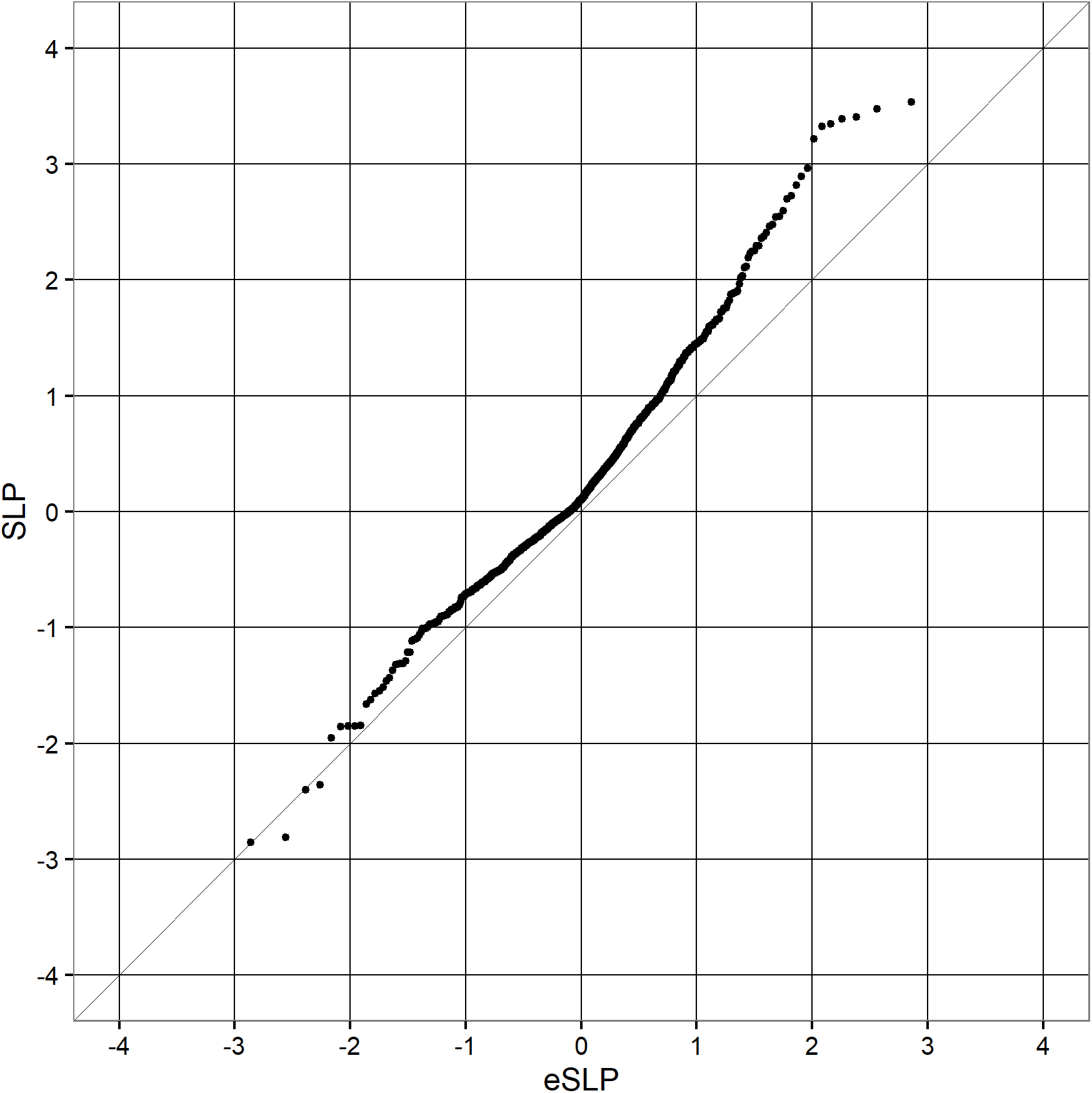
QQ plot for set-wise SLPs obtained when including all covariates for GO gene sets against the expected SLP if all sets were non-overlapping and independent.

## Discussion

This study demonstrates that it possible to use a logistic regression approach to carry out a weighted burden analysis, including all variants in a gene weighted according to function and rarity, while including other important covariates. The example analysis produces results which are broadly similar to previous analyses of this dataset, although with less strong support for the previously implicated gene sets.

An important finding is that including the population principal components is able to completely control for the effect of having an excess of cases with a strong Finnish ancestry component. This means that the complete dataset can be analysed if desired. Of course, there might be arguments against doing this. If different causal variants are present in different populations then analysing an admixed population might reduce power but in general it seems advantageous to be able to incorporate ancestry as a covariate. From a theoretical point of view we might expect that incorporating known risk factors as covariates could increase the power to detect new associations. In practice, including the PRS and pathogenic CNVs and sequence variants has only a small effect on the results obtained in the present example.

Although the main emphasis of the study is simply to demonstrate the feasibility of the approach, it is possible to speculate on why it failed to provide any new, interesting results. One problem may well be that the default scheme for weighting variants according to predicted functional effects is sub-optimal. Many criticisms of it could be made, for example that LOF variants and 5’UTR variants should be given higher weights. For the future, it would be worthwhile to make attempts to produce a weighting scheme which was informed by the increasing empirical knowledge becoming available about the kinds of genetic variation most likely to have major effects on phenotype. Another approach would be to attempt to fit the weights directly and this could easily be done within the context of logistic regression analysis if it were thought that the dataset was informative enough. A similar approach has been applied in the context of generating an exome-wide genetic risk score^22^.

Incorporating pathogenic variants had little effect on the results obtained and one reason for this is that only 1% of subjects were identified as possessing such a variant.

It would be reasonable to hope that as further studies discover additional variants to be pathogenic so the proportion of subjects possessing one them will increase and they will have a larger impact on the results of multivariate analyses. However another factor to consider is that there were almost certainly errors in the CNV calls and that had the calls been accurate then they might have had more influence on the results. This is simply a feature of using a downloaded VCF file to provide an example dataset.

In a real-world association study using whole exome sequence data one would have the raw reads available and could make more reliable calls from those, and of course one might also carry out comparative genomic hybridisation analysis to obtain accurate calls.

Although the method described here does not seem to have marked benefits for this dataset, it does offer an approach that will be useful to apply as additional risk factors, both genetic and non-genetic, are characterised and need to be incorporated into analyses. An obvious example would be for a disease such as ischaemic heart disease (IHD), where rare and common genetic variants along with environmental risk factors all make substantial contributions^23^. The application of logistic regression has been described as a way to incorporate information from covariates into association studies seeking to identify variants and genes associated with disease. However the same framework could then readily be translated into a way to characterise risk of disease for an individual. One could fit a model incorporating relevant risk factors using a study population and then take the fitted coefficients for that model and apply them to a genotyped individual subject to obtain an estimate of their level of risk. It has already been proposed that the PRS IHD for could have clinical utility in deciding whether to prescribe statins for a patient^24^. Using a model such as the one proposed would allow an overall risk score to be produced from a single combined analysis including factors such as age, gender, smoking status, ancestry, IHD PRS and rare coding variants. One can envisage in future that one might even be able to partition overall risk in clinically significant ways, for example with a contribution related to dyslipidemia, a contribution related to clotting and a contribution related to hypertension. This could allow the subclassification of patients to support personalised approaches to treatment interventions.

## Software availability

The code and documentation for SCOREASSOC, PATHWAYASSOC and GENEVARASSOC is available from https://github.com/davenomiddlenamecurtis/scoreassoc and https://github.com/davenomiddlenamecurtis/geneVarAssoc.

## Acknowledgements

The datasets used for the analysis described in this manuscript were obtained from dbGaP at http://www.ncbi.nlm.nih.gov/gap through dbGaP accession number phs000473.v2.p2. Samples used for data analysis were provided by the Swedish Cohort Collection supported by the NIMH grant R01MH077139, the Sylvan C. Herman Foundation, the Stanley Medical Research Institute and The Swedish Research Council (grants 2009-4959 and 2011-4659). Support for the exome sequencing was provided by the NIMH Grand Opportunity grant RCMH089905, the Sylvan C. Herman Foundation, a grant from the Stanley Medical Research Institute and multiple gifts to the Stanley Center for Psychiatric Research at the Broad Institute of MIT and Harvard.

## References

1 Morris AP, Zeggini E. An evaluation of statistical approaches to rare variant analysis in genetic association studies. Genet Epidemiol 2010; 34: 188–193.

2 Purcell SM, Moran JL, Fromer M et al. A polygenic burden of rare disruptive mutations in schizophrenia. Nature 2014; 506: 185–90.

3 Madsen BE, Browning SR. A groupwise association test for rare mutations using a weighted sum statistic. PLoS Genet 2009; 5: e1000384.

4 Curtis D. A rapid method for combined analysis of common and rare variants at the level of a region, gene, or pathway. Adv Appl Bioinform Chem 2012; 5: 1–9.

5 Curtis D. Pathway analysis of whole exome sequence data provides further support for the involvement of histone modification in the aetiology of schizophrenia. Psychiatr Genet 2016; 26: 223–7.

6 Marshall CR, Howrigan DP, Merico D et al. Contribution of copy number variants to schizophrenia from a genome-wide study of 41,321 subjects. Nat Genet 2016; 49: 27–35.

7 Singh T, Kurki M, Curtis D, Purcell S, et al. Rare SETD1A loss-of-function variants are associated with schizophrenia and developmental disorders. Nat Neurosci 2016.

8 Steinberg S, Gudmundsdottir S, Sveinbjornsson G et al. Truncating mutations in RBM12 are associated with psychosis. Nat Genet 2017. doi:10.1038/ng.3894.

9 Genovese G, Fromer M, Stahl EA et al. Increased burden of ultra-rare protein-altering variants among 4,877 individuals with schizophrenia. Nat Neurosci 2016; 19: 1433–1441.

10 Curtis D, Coelewij L, Liu S-H, Humphrey J, Mott R. Weighted Burden Analysis of Exome-Sequenced Case-Control Sample Implicates Synaptic Genes in Schizophrenia Aetiology. Behav Genet 2018;: 203521.

11 Schizophrenia Working Group of the Psychiatric Genomics Consortium. Biological insights from 108 schizophrenia-associated genetic loci. Nature 2014; 511: 421–427.

12 McLaren W, Gil L, Hunt SE et al. The Ensembl Variant Effect Predictor. Genome Biol 2016; 17: 122.

13 Adzhubei I, Jordan DM, Sunyaev SR. Predicting functional effect of human missense mutations using PolyPhen-2. Curr Protoc Hum Genet 2013; 7 **Unit7.20**. doi:10.1002/0471142905.hg0720s76.

14 Kumar P, Henikoff S, Ng PC. Predicting the effects of coding non-synonymous variants on protein function using the SIFT algorithm. Nat Protoc 2009; 4: 1073–1081.

15 King D. Dlib-ml: A Machine Learning Toolkit. J Mach Learn Res 2009; 10: 1755–1758.

16 Purcell S, Neale B, Todd-Brown K et al. PLINK: a tool set for whole-genome association and population-based linkage analyses. Am J Hum Genet 2007; 81: 559–75.

17 Chang CC, Chow CC, Tellier LC, Vattikuti S, Purcell SM, Lee JJ. Second-generation PLINK: rising to the challenge of larger and richer datasets. Gigascience 2015; 4: 7.

18 Purcell SM, Wray NR, Stone JL et al. Common polygenic variation contributes to risk of schizophrenia and bipolar disorder. Nature 2009; 10: 8192–8192.

19 Rujescu D, Ingason A, Cichon S et al. Disruption of the neurexin 1 gene is associated with schizophrenia. Hum Mol Genet 2009; 18: 988–996.

20 Ching MSL, Shen Y, Tan W-H et al. Deletions of NRXN1 (neurexin-1) predispose to a wide spectrum of developmental disorders. Am J Med Genet B Neuropsychiatr Genet 2010; 153B: 937–47.

21 R Core Team. R: A language and environment for statistical computing. R Foundation for Statistical Computing: Vienna, Austria., Austria., 2014.

22 Curtis D. Construction of an exome-wide risk score for schizophrenia based on a weighted burden test. Ann Hum Genet 2017.

23 Knowles JW, Ashley EA. Cardiovascular disease: The rise of the genetic risk score. PLOS Med 2018; 15: e1002546.

24 Knowles JW, Ashley EA. Cardiovascular disease: The rise of the genetic risk score. PLOS Med 2018; 15: e1002546.

25 Hamosh A, Scott AF, Amberger JS, Bocchini CA, McKusick VA. Online Mendelian Inheritance in Man (OMIM), a knowledgebase of human genes and genetic disorders. Nucleic Acids Res 2005; 33: D514–7.

26 Fagerberg L, Hallstrom BM, Oksvold P et al. Analysis of the Human Tissue-specific Expression by Genome-wide Integration of Transcriptomics and Antibody-based Proteomics. Mol Cell Proteomics 2014; 13: 397–406.

27 Wagnon JL, Briese M, Sun W et al. CELF4 Regulates Translation and Local Abundance of a Vast Set of mRNAs, Including Genes Associated with Regulation of Synaptic Function. PLoS Genet 2012; 8: e1003067.

28 Samocha KE, Robinson EB, Sanders SJ et al. A framework for the interpretation of de novo mutation in human disease. Nat Genet 2014; 46: 944–950.

29 Deciphering Developmental Disorders Study. Prevalence and architecture of de novo mutations in developmental disorders. Nature 2017; 542: 433–438.

30 Fromer M, Pocklington AJ, Kavanagh DH et al. De novo mutations in schizophrenia implicate synaptic networks. Nature 2014; 506: 179–184.

31 Kirov G, Pocklington AJ, Holmans P et al. De novo CNV analysis implicates specific abnormalities of postsynaptic signalling complexes in the pathogenesis of schizophrenia. Mol Psychiatry 2012; 17: 142–153.

32 Darnell JC, Van Driesche SJ, Zhang C et al. FMRP Stalls Ribosomal Translocation on mRNAs Linked to Synaptic Function and Autism. Cell 2011; 146: 247–261.

33 Robinson EB, Neale BM, Hyman SE. Genetic research in autism spectrum disorders. Curr Opin Pediatr 2015; 27: 685–691.

34 Cahoy JD, Emery B, Kaushal A et al. A Transcriptome Database for Astrocytes, Neurons, and Oligodendrocytes: A New Resource for Understanding Brain Development and Function. J Neurosci 2008; 28: 264–278.

35 Lek M, Karczewski KJ, Minikel E V et al. Analysis of protein-coding genetic variation in 60,706 humans. Nature 2016; 536: 285–291.

36 Bayés A, van de Lagemaat LN, Collins MO et al. Characterization of the proteome, diseases and evolution of the human postsynaptic density. Nat Neurosci 2011; 14: 19–21.

37 Weyn-Vanhentenryck SM, Mele A, Yan Q et al. HITS-CLIP and Integrative Modeling Define the Rbfox Splicing-Regulatory Network Linked to Brain Development and Autism. Cell Rep 2014; 6: 1139–1152.

38 Pirooznia M, Wang T, Avramopoulos D et al. SynaptomeDB: an ontology-based knowledgebase for synaptic genes. Bioinformatics 2012; 28: 897–9.

39 Cotton AM, Ge B, Light N, Adoue V, Pastinen T, Brown CJ. Analysis of expressed SNPs identifies variable extents of expression from the human inactive X chromosome. Genome Biol 2013; 14: R122.

40 Moeschler JB. Genetic Evaluation of Intellectual Disabilities. Semin Pediatr Neurol 2008; 15: 2–9.

41 Gécz J, Shoubridge C, Corbett M. The genetic landscape of intellectual disability arising from chromosome X. Trends Genet 2009; 25: 308–316.

42 Moeschler JB, Shevell M, American Academy of Pediatrics Committee on Genetics. Clinical Genetic Evaluation of the Child With Mental Retardation or Developmental Delays. 2006; 117: 2304–2316.

43 Rauch A, Hoyer J, Guth S et al. Diagnostic yield of various genetic approaches in patients with unexplained developmental delay or mental retardation. Am J Med Genet Part A 2006; 140A: 2063–2074.

